# ITNR: Inversion Transformer-based Neural Ranking for Cancer Drug Recommendations

**DOI:** 10.1101/2023.03.16.533057

**Authors:** Shahabeddin Sotudian, Ioannis Ch. Paschalidis

**Author notes:** Corresponding author, (S. Sotudian); (.I.Ch. Paschalidis).

## Abstract

Personalized drug response prediction is an approach for tailoring effective therapeutic strategies for patients based on their tumors’ genomic characterization. The current study introduces a new listwise Learning-to-rank (LTR) model called Inversion Transformer-based Neural Ranking (ITNR). ITNR utilizes genomic features and a transformer architecture to decipher functional relationships and construct models that can predict patient-specific drug responses. Our experiments were conducted on three major drug response data sets, showing that ITNR reliably and consistently outperforms state-of-the-art LTR models.

**Highlights:**

- The proposed framework is a transformer-based model to predict drug responses using RNAseq gene expression profile, drug descriptors and drug fingerprints.
- ITNR utilizes a Context-Aware-Transformer architecture as its scoring function that ensures the modeling of inter-item dependencies.
- We introduced a novel loss function using the concept of Inversion and Approximate Permutation matrices.
- Our computational results indicated that our method leads to substantially improved performance when compared to the baseline methods across all performance metrics, which can lead to selecting highly effective personalized treatment.

## 1. Introduction

Conventional Machine Learning (ML) algorithms are generally designed to minimize regression or classification errors [12, 13, 16, 32, 42]. Many real-world applications, on the other hand, deemphasize prediction accuracy and prioritize the correct ordering among all the instances [33, 34]. The aim of ***Learning-to-rank (LTR)*** methods is to apply ML techniques to solve ranking problems and predict the optimal ordering among the instances according to their degrees of relevance, importance, or preference as defined in the specific application.

Ranking plays a central role in a wide variety of applications, including document retrieval, online advertisement, drug discovery, machine translation, feature selection, document summarization, definition search, question answering, and recommendation systems, among others [27, 25]. In these applications, it is highly desirable to design a model that places relevant/important items at the top of the ranking list. In this work, and without loss of generality, we focus on cancer ***Drug Response Prediction (DRP)*** where the goal is to prescribe an optimal therapeutic option for each patient based on their cancer’s unique molecular fingerprints (i.e., the goal of “precision oncology”). Not every patient responds to medical treatment in the same way. Effectiveness of a medication is influenced by various factors including physiological, pathological, environmental, and genetic factors [40]. Since in-vitro experiments are exceedingly costly and time-consuming, DRP algorithms could serve as promising strategies for the accurate prediction of optimal drug therapies based upon the personalized molecular profiles of patient tumors [35].

The confluence of efficient computational tools and a significant number of samples has ushered in a new generation of ML models for drug recommendation. The literature offers a variety of traditional approaches from Support Vector Machines (SVMs); Principal Component Regression; Ridge, LASSO, and Elastic Net Regressions; to more advanced methods such as multiple-output [33], multiple-kernel [19], and multiple-task learning [30] techniques. As for traditional methods, comprehensive comparative studies [15, 35] demonstrated that Elastic Net or Ridge regression-based models will most likely yield the most accurate predictors. On the other hand, deep learning (DL) models have demonstrated their superiority in capturing the non-linear and complex relationships of biological data better than the traditional algorithms [1]. Representative DL-based algorithms include [2, 29, 21]. These works, [5, 1] and [40] provided a comprehensive analysis of DRP models and other related topics such as data integration, feature selection, experimental settings, combination therapy, and so on.

In the current work, we seek to use an under-explored approach for drug response prediction problems, namely ***transformer*** models. The application of transformer-based techniques in LTR signifies the high capability of these models in various applications [21]. Equipped with this perspective, we make the following contributions. We developed our LTR framework using a context-aware scoring function and an inversion-based loss function. Unlike the majority of LTR models whose scoring functions score items separately, transformer models (i.e., a popular self-attention-based neural machine translation architecture) allow for the modeling of inter-item dependencies. We adopt the Context-Aware-Transformer [21] which is a special case of the encoder part of the transformer. Additionally, this architecture takes into account the inter-dependency of scores between items in the computation of items’ scores. We also proposed a loss function using the concept of *Inversion* and *Approximate Permutation* matrices.

In experiments, our framework yields state-of-the-art results in three major drug response data sets, showing that our model maintains a consistently good performance under various experimental settings. Thus, the proposed architecture can capture local context information and cross-item interactions that lead to a reliable drug recommendation system.

### Notational conventions

We use boldfaced lowercase letters to denote vectors, ordinary lowercase letters to denote scalars, boldfaced uppercase letters to denote matrices, and calligraphic capital letters to denote sets. All vectors are column vectors. For space saving reasons, we write **x** to denote the column vector (*x*_1_…, *x*_dim(X)_), where dim(**x**) is the dimension of **x**. We use prime to denote the transpose, and 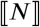 for the set {1,…, *N*} for any integer *N*.

## 2. Materials and Methods

### 2.1. Problem Formulation

Data in a drug ranking problem consist of a set of triples (cell line, drug, drug response score). A cell line-drug pair is represented by a feature vector. Our ultimate goal is to select the most effective drugs from a set of drugs based on their response. We characterize a drug ranking data set with tuples 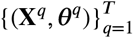 where 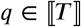 indexes cell lines, and **X**^*q*^ and ***θ**^q^* represent the list of cell line-drug pairs and corresponding drug response scores, respectively. For the *q*-th cell line, we have *n_q_* drugs. 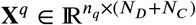 has rows 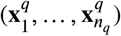, each of which is a (*N_D_* + *N_C_*)-dimensional cell line-drug vector, formed as the concatenation of an (*N_C_*)-dimensional cell line feature vector (i.e., gene expression for cell lines) and an (*N_D_*)-dimensional drug feature vector. The vector 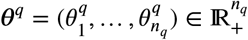 contains the corresponding ground-truth drug response scores. The drug response scores are in [0, 1] and a higher score (in our data) implies a more effective drug. Our goal is to learn a scoring function 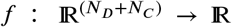 from a training data set with given drug response scores that minimizes the empirical loss:

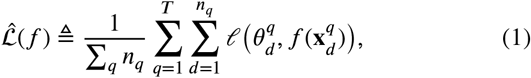

where 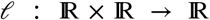 is a loss function. For a new cell line-drug matrix 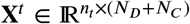, we can obtain the predicted ranking list by ranking the rows in **X**^*t*^ based on their inferred ranking scores 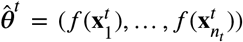. Two pivotal elements of an LTR framework are a scoring function and a loss function. In subsequent subsections, we describe the construction of our LTR framework using a context-aware scoring function and an inversion-based loss function.

### 2.2. Multi-Headed Self-Attention Scoring Function

Most algorithms in the DRP literature are trained to optimize a loss function that may not capture the interactions among drugs [35]. Moreover, they score them individually at inference-time without regard to any *mutual influences* among the drugs. Given that context-aware models based on transformers [21] have been successfully used for document retrieval, we adopt a similar approach of applying a context-aware ranker for our scoring function. The transformer architecture proposed by [21] plays the role of the ***Multi-Headed Self-Attention (MHSA)*** scoring function in our LTR framework. The self-attention mechanism of this architecture aims to handle long-range and inter-item dependencies. Since the MHSA scoring function scores a drug by considering all other drugs applicable to a cell line, it fully captures the interactions among drugs.

We consider the list’s items as our tokens where item features are token embeddings. We feed these embeddings to an *Encoder Layer.* The attention mechanism is the core of the encoder layer to catalyze learning the higher-order representations of items in the list. Mapping query (**q**_*i*_), key (**k**_*i*_), and value (**v**_i_) vectors to a higher level representation by taking a weighted sum of the values over all items is the essence of the self-attention mechanism. We use the *Scaled Dot-Product* form to compute attention as follows:

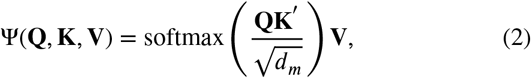

where **Q** is a query matrix that contains all items (queries) in the list; **K** and **V** are the key and value matrices, respectively. To empower the model to leverage the order of the input tokens, we use fixed positional encodings for the input embeddings as follows:

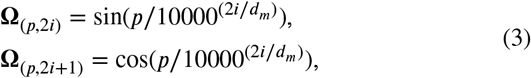

where *p* refers to the position, and *i* is the dimension. Here, **Ω** is a matrix and positional encoding is a system to encode each position into a vector. For instance, given that the model uses 256 dimensions for positional encoding (*d_m_* = 256), we represent each element of the feature vector (i.e., token) as a 256-dimensional vector. In the display above, *p* is an integer from 0 to a pre-defined maximum number of tokens minus 1. Since we have 256 dimensions, we can define 128 pairs of sine and cosine values. Accordingly, the value of *i* goes from 0 to 127. By ensembling multiple attention modules (a.k.a Multi-Head Attention (MHA)), we can improve the ability of the model to learn representations from various subspaces of the input data. MHA can be expressed as:

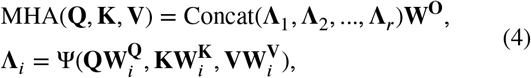

where in the above equations, each head **Λ**_*i*_ (out of *r* heads) refers to the *i*-th attention mechanism of Equation (2), and 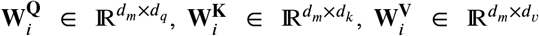 and 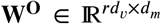 are learnable matrices. Moreover, *r* refers to the number of parallel attention layers or heads. Here, Concat(·) represents a concatenation operation. It is extremely advantageous to perform the self-attention operation several times and concatenate the outputs. The main problem is the growing size of the resulting output vector. This can be solved by linearly projecting the matrices **Q**, **K**, and **V** to *d_q_*, *d_k_*, and *d_ν_*-dimensional spaces *r* times, respectively. Note that we typically use *d_q_* = *d_k_* = *d_ν_* = *d_m_*/*r*. We refer the interested reader to [37] for more information.

In our transformer model, we use a more complex architecture by stacking multiple encoder blocks. The main components of an encoder block including an MHA layer with a skip connection, layer normalization, time-distributed feed-forward layer, and dropout layer can be seen in Figure 1. For more information regarding these components, please refer to [37]. Specifically, the encoder part of the Transformer includes *N* encoder blocks, *H* heads, and hidden dimension *d_h_*. First, a shared fully connected (FC) input layer of size *d_fc_* is applied to each item. Then, we feed hidden representations to the encoder part of the transformer. Our final scoring model can be achieved by iteratively stacking multiple encoder blocks where the output of each block is fed into the next one. The final model can be expressed as follows:

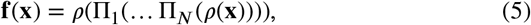

where *ρ*(·) represents a projection onto a fully-connected layer and ⊓_*i*_(·) refers to a single encoder block. We define a single encoder block ⊓·(·) as:

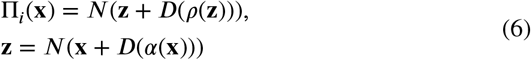

where *α*(·) is the MHA module, *N*(·) refers to the layer normalisation, and *D*(·) is the dropout layer. Eventually, a score for each item is calculated using a shared fully-connected layer. Figure 1 provides a schematic overview of our LTR framework.

**Figure 1:**
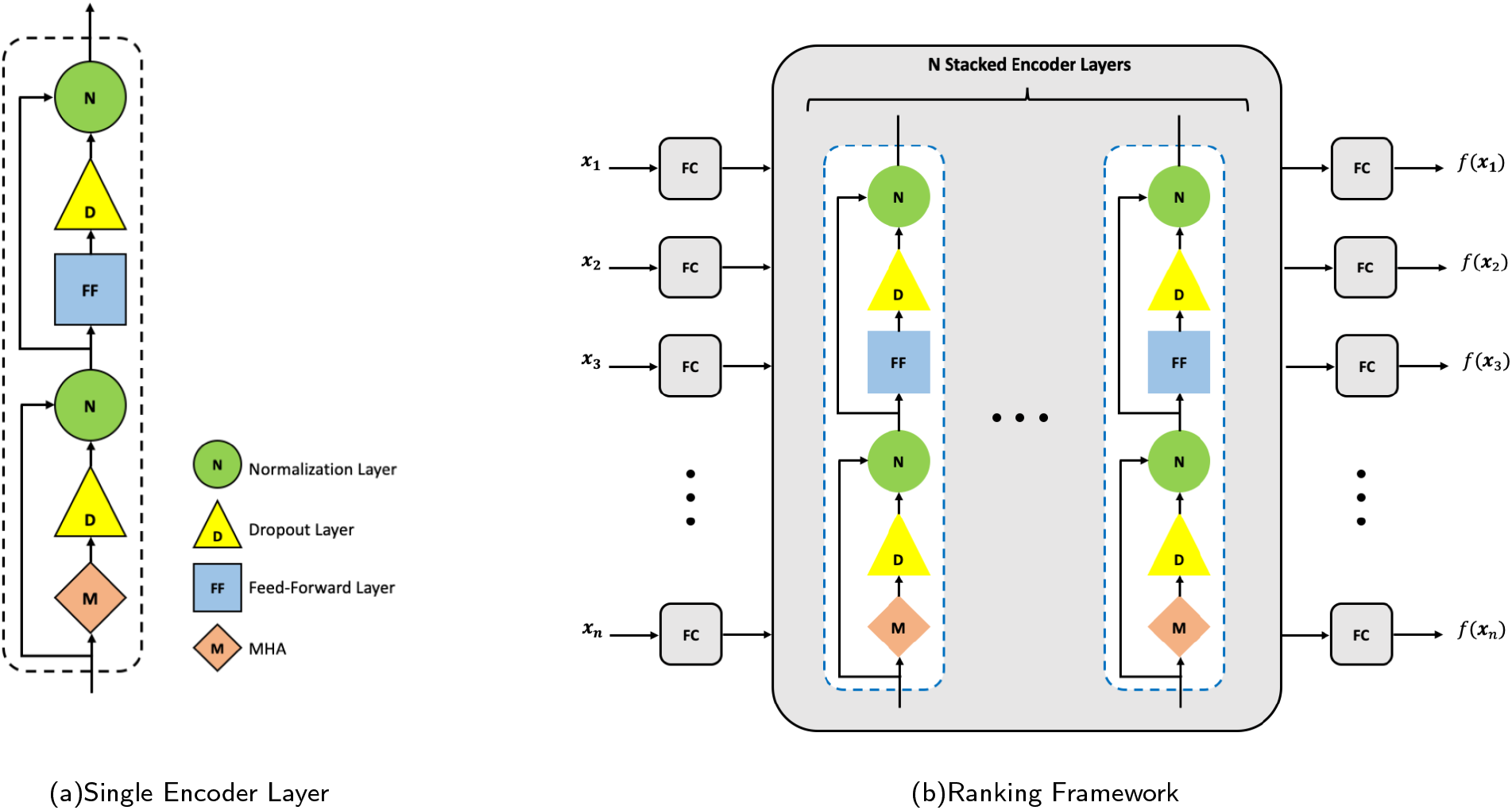
Architecture for the context-aware ranking model. After feeding an item’s features into a fully connected layer, we pass its output through *N* encoders blocks. Finally, another fully-connected layer is used to calculate scores.

### 2.3. Inv-Rank Loss Function

In the previous subsection, we presented the scoring function of our ranking framework. The architecture of this model allows the networks to exploit local features. More importantly, the final score for each item will be calculated by considering all other items on the list. Now, we can use the scores and the ground truth labels to optimize any desired ranking loss.

It has been demonstrated that the sorting operator can be approximated by the induced permutation matrix, *P*_*sort*(S)_ [21]. We use the concept of *permutation matrices* to define our proposed loss function. Permutation matrices are both **doubly-stochastic** (i.e., a square matrix with entries in [0, 1] where every row and column sum to one) and **unimodal** (i.e., a square matrix with entries in [0, 1] where each row sums to one, but also has the constraint that the maximizing entry in every row should have a unique column index) [10, 21]. Assume **A_s_** represents the matrix of absolute pairwise score differences of **s** with the *i, j*-th element given by **A_s_**[*i, j*] = |*s_i_* – *S_j_*|. Grover et al. [10] proposed a continuous relaxation of the permutation matrix 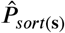 in the space of unimodal row-stochastic matrices. The *i*-th row of 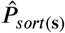 can be computed as follows:

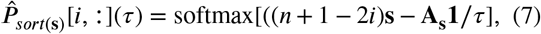

where **1** is an all-one vector, *n* is the number of items in a list, and *τ* behaves like a temperature knob that controls the degree of approximation and the variance of the gradients (i.e., lower *τ* leads to better approximation and higher variance). Moreover, softmax is a function that scales the values in the list and transforms them into values between 0 and 1 such that all values in the returned list sum to 1 (i.e., they can be interpreted as probabilities). Essentially, if we left-multiply a column vector of scores by its 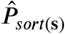, we can achieve the approximated sorted list [22]. Note that we perform Sinkhorn scaling [31] to obtain *doubly stochastic* permutation matrices. The Sinkhorn scaling method normalizes all rows and columns; this process is repeated until convergence is achieved (i.e., 30 iterations or the greatest gap between the sum of a row or column and one must be maintained below 10^-6^, whichever occurs first). Assume we use the MHSA scoring function to calculate scores **s**^*q*^ = *f*(**X**^*q*^) for the *q*-th cell line **X**^*q*^. We also have the ground truth labels *θ^q^* for the cell line. We define true and predicted *Weighted Approximate Permutation (WAP) matrices* using Equation (7) as:

- 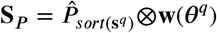: predicted WAP matrix induced by the scores **s**^*q*^,
- 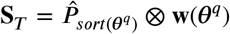: true WAP matrix induced by the ground truth labels ***θ***^*q*^,

where 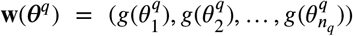 is the drug importance vector, and *g*(*x*) = 2^*x*^ – 1 is a famous LTR gain function. Here, ⊗ indicates the multiplication of columns of 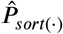 by elements of the vector **w**(·) (i.e., multiply the first column by the first element, second column by the second element and so on). The WAP matrix (i.e., **S**_*P*_ and **S**_*T*_) captures the relative position of a drug in a list. Each column of this matrix refers to a drug and the non-zero element of a column represents the predicted/true position of a drug in a ranking list. Note that if the ground truth labels of multiple drugs in a list are the same, then we have multiple ideal positions for each of them. Thus, multiple elements of those columns (i.e., drugs) will be non-zero which shows the potential predicted/true positions of those drugs in the list. We use **w** to force our model to focus on more sensitive drugs (i.e., it gives higher weights to more sensitive drugs) rather than insensitive drugs. Consequently, the model pushes sensitive drugs to the top of the ranking list.

We use the concept of *Inversion* to define our loss function. Inversion can be defined as a pair of elements that are out of their correct order in a permutation. Let *π* be a permutation. If *i* < *j* and *π*(*i*) > *π*(*j*), either the pair of places (*i, j*) or the pair of elements (*π*(*i*), *π*(*j*)) is called an inversion of *π*. We define **Θ** to capture inversions in a permutation matrix as follows:

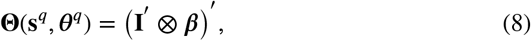

where 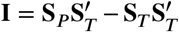 and ***ß*** = (1, 2, 3,…, *n_q_*). Precisely, **I** measures the deviation between predicted and true WAPs and penalizes the model for any inversion. The rows of a WAP matrix represent ranks. There exist various degrees of inversion. For instance, there is a big difference between (*π*(1), *π*(3)) and (*π*(1), *π*(7)). Our model should penalize (*π*(1), *π*(7)) more than (*π*(1), *π*(3)). To that end, we multiply the rows of the **I** matrix by the elements of ***β***. Therefore, our model places greater emphasis on high inversions. Eventually, we define the **Inv-Rank** loss of all cell lines in our training data set as follows:

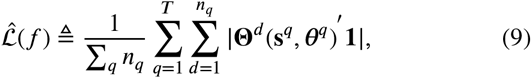

where **Θ**^*d*^ (**s**^*q*^, **θ**^*q*^) is the *d*-th column of **Θ**(**s**^*q*^, **θ**^*q*^). Since **S**_*P*_ is differentiable with respect to the elements of **s** [10], it is easy to show that the proposed loss function is a differentiable function of scores. Therefore, the SGD method can be used to optimize the loss function.

### 2.4. Data sets and Pre-processing Steps

The current study focused on single-drug response prediction and we designed, trained, and evaluated all models using the cell line data and drug sensitivity data from the Predictive Oncology Model & Data Clearinghouse hosted at the National Cancer Institute [40]. We used three main drug-cell line data sets from the *Cancer Cell Line Encyclopedia (CCLE),* the *Genomics of Drug Sensitivity in Cancer (GDSC),* and the *Genentech Cell Line Screening Initiative (gCSI)* studies; and molecular descriptors generated using the Dragon 7.0 and the Mordred software packages [40]. Specifically, we represented a cell line using its RNAseq gene expression profile [36]; and we used drug descriptors and drug fingerprints to characterize a drug. The Area Under the drug response Curve (AUC) was used to quantify drug sensitivity. AUC ∈ [0, 1] can be compared across studies and a lower value indicates higher drug sensitivity. We modified drug sensitivities so that a higher value indicates a more effective drug (i.e., one minus AUC). The details of the data sets can be found in [40]. To follow the best practice of LTR [25], the continuous drug responses were converted to graded ones. Accordingly, we classified drugs into three categories 0 (i.e., “insensitive”), 1 (i.e., “sensitive”), and 2 (i.e., “highly sensitive”). For the *k*-th cell line, let *P*_80_ and *P*_90_ denote the 80-th and 90-th percentiles of its drug response values [*r*_1*k*_, *r*_2*k*_,…, *r_N_D_k_*}, respectively. Then, the drug relevance score for the *i*-th drug, 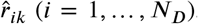, can be computed as:

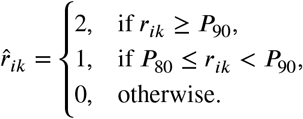

We also performed several pre-processing steps to reduce the training complexity and improve the overall performance. We standardize the features by subtracting the mean and scaling to unit variance [28]. Originally, RNAseq gene expressions are represented by approximately 17,000 features. Several studies [6] demonstrated that the LINCS1000 gene set [17] can outperform or achieve similar performance compared to any other superset of LINCS1000. Therefore, we only included LINCS1000 genes in our analysis. Moreover, the data sets include 3838 molecular descriptors and 1024 path fingerprint features to represent a drug. To further reduce the dimensionality of our data and select informative features, we used a regression model as the feature selection method. Specifically, we considered drug-cell line vectors (i.e., 5862 features including 1000 genes, 3838 molecular descriptors, and 1024 path fingerprint features) and the response values (i.e., one minus AUCs) as independent and dependent variables, respectively. First, the regression model is applied to these variables and the importance of each feature is obtained through regression coefficients. After standardizing the variables, a larger absolute regression coefficient indicates that this specific variable has more influence on drug sensitivity (i.e., dependent variable). We have two groups of features, namely 1000 cell line features and 4862 drug features. We kept 500 genes out of 1000 gene features with the highest coefficient values. Similarly, we selected 500 features out of the 4862 drug features. Consequently, we used 1000 selected gene-drug features in our experiments. We repeated this procedure for all three data sets.

### 2.5. Performance metrics

Two main LTR evaluation metrics, namely NDCG@k and MRR@k are used to assess the performance of the models. Let *D*(*s*) = 1/log(1 + *s*) be a discount function, *g*(*s*) = 2^*s*^ – 1, a monotonically increasing gain function, and 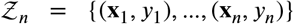 a set of items ordered according to their ground-truth labels, with **x**_*i*_ and *y_i_* being an item feature vector and score, respectively. Moreover, let 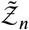 be a ranked list for 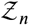 according to the **y** scores. We define the *Discounted Cumulative Gain (DCG)* of 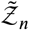 as 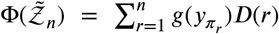, where *π_r_* is the index of the item ranked at position *r* of 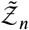. The Ideal DCG (IDCG), 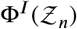 is the DCG score of the ideal ranking result. NDCG normalizes DCG by the IDCG and can be calculated by 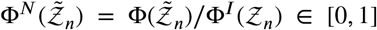. To force the metric to focus on the top-k items, we use NDCG@k, which is the top-k version of NDCG, where the discount function is *D*(*s*) = 0 for *s* > *k*. Mean Reciprocal Rank (MRR) puts a high focus on the most relevant item of a list. Assume *r_i_* denotes the rank of the most relevant item in the *i*-th list, then the reciprocal rank is defined as *R_i_* = 1/*r_i_*. For *N* lists, the MRR is the mean of the N reciprocal ranks, 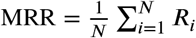. MRR@k is simply the top-k version of MRR. The significant difference between MRR and NDCG is that NDCG distinguishes between “partially sensitive” and “highly sensitive” drugs while MRR only focuses on the most sensitive drugs. From now on, we use NDCG@k and MRR@k to denote the mean NDCG@k and the mean MRR@k (i.e., the mean of the performance metric for all lists in our test set).

### 2.6. Experimental settings and hyper-parameter optimization

We conducted our experiments based on the standard supervised LTR framework [20]. The authors of LETOR [25] partitioned the LTR data sets into five parts for five-fold cross-validation where three parts were used for training, one part for validation (i.e., tuning the hyperparameters of the learning algorithms), and the remaining part for evaluating the performance of the learned model. We followed the same procedure, partitioning our data sets into five-folds, and conducting five-fold cross-validation to train the models. The hyperparameters were tuned on the validation sets (i.e., we optimized the hyperparameters to maximize the NDCG score) and the average on the test sets over the 5 folds was reported in the various tables. For more details on the parameter-tuning procedure and experimental settings, please refer to Appendix.

### 2.7. Competing Methods

We compared our proposed ranking algorithm with three types of algorithms, namely Transformer-based Neural Ranking (TNR), Deep Neural Networks (DNN), and traditional models. In recent years, the attention mechanism in transformers began a revolution in deep neural networks that led to major advances in the performance of many models obtained in this fashion. We compared our model with four TNR models (i.e., NDCGLoss2++[38], ListMLE [41], ApproxNDCG [24], and RankNet [4]) to ensure its superior performance and stability compared to similar methodologies. Second, we also compared ITNR with other deep learning-based models. To that end, we compared ITNR with the so-called DNN-Sakellaropoulos (DNN-S) [29]. DNN-S has been reported as one of the best DNN models in the DRP literature [18, 35]. Third, several DRP comparative studies [11, 8, 26] reported tree-based models as one of the best-performing algorithms for drug recommendation. Specifically, LambdaMART_MAP_ [39] has been shown repeatedly to surpass other LTR methods such as Coordinate Ascent, Random Forests, BoltzRank, RankBoost, AdaRank, SoftRank, and so on [8, 3, 38]. Finally, we compared ITNR with Elastic Net Regression (ENR) as a traditional DRP model. Multiple studies [14, 35, 9] have suggested that elastic net will most likely yield the most accurate predictors for drug response prediction.

## 3. Experimental Results

In Table 1, we summarized the performance of the models on various DRP data sets. We report NDCG@k and MRR@k, both computed out-of-sample (i.e., test set not used for training the model). The average on the test set over the 5 folds was reported. Bold and underlined numbers indicate the best performance among all methods for each metric. Bold numbers demonstrate the second-best performance among all methods for each metric.

**Table 1.** Performance Comparison of Ranking Methods on CCLE, GDSC, and gCSI Data Sets.

| Algorithms | NDCG@5 | NDCG@10 | NDCG@25 | MRR@5 | MRR@10 | MRR@25 |
| --- | --- | --- | --- | --- | --- | --- |
| **CCLE Data set** |  |  |  |  |  |  |
| ITNR | **92.15%** | **94.03%** | **94.09%** | **93.87%** | **93.90%** | **93.90%** |
| TNR-NDCGLoss2++ | 91.87% | 93.71% | 93.85% | 92.93% | 92.96% | 92.96% |
| TNR-ListMLE | 91.78% | 93.41% | 93.93% | 93.62% | 93.71% | 93.71% |
| TNR-RankNet | **92.39%** | 93.76% | **94.00%** | 93.52% | 93.52% | 93.52% |
| TNR-ApproxNDCG | 90.71% | 92.40% | 92.49% | 88.05% | 88.08% | 88.08% |
| DNN-S | 92.06% | 93.63% | 93.99% | **93.71%** | **93.74%** | **93.76%** |
| LambdaMART<sub>MAP</sub> | 92.03% | 93.46% | 93.72% | 92.25% | 92.28% | 92.28% |
| ENR | 91.73% | **93.85%** | 93.94% | 93.55% | 93.64% | 93.64% |
| **GDSC Data set** |  |  |  |  |  |  |
| ITNR | **83.36%** | **79.26%** | **82.40%** | **92.74%** | **92.76%** | **92.76%** |
| TNR-NDCGLoss2++ | 82.01% | 78.47% | 82.12% | 91.33% | 91.33% | 91.33% |
| TNR-ListMLE | 80.89% | 77.08% | 80.04% | 88.77% | 88.79% | 88.79% |
| TNR-RankNet | **82.69%** | **78.59%** | 82.38% | **92.16%** | **92.16%** | **92.16%** |
| TNR-ApproxNDCG | 82.22% | 77.84% | 80.86% | 90.79% | 90.79% | 90.79% |
| DNN-S | 77.80% | 76.18% | 80.15% | 83.96% | 83.96% | 83.96% |
| LambdaMART<sub>MAP</sub> | 81.31% | 78.24% | **82.79%** | 89.62% | 89.64% | 89.64% |
| ENR | 76.97% | 74.89% | 78.39% | 83.70% | 83.82% | 83.82% |
| **gCSI Data set** |  |  |  |  |  |  |
| ITNR | **80.36%** | **83.20%** | **83.35%** | **77.67%** | **77.71%** | **77.71%** |
| TNR-NDCGLoss2++ | 78.91% | 82.62% | 82.70% | 76.38% | 76.52% | 76.52% |
| TNR-ListMLE | 76.97% | 81.08% | 81.31% | 75.38% | 75.47% | 75.47% |
| TNR-RankNet | 79.59% | **82.95%** | **82.99%** | 76.33% | 76.47% | 76.47% |
| TNR-ApproxNDCG | 78.56% | 82.50% | 82.61% | 76.09% | 76.09% | 76.09% |
| DNN-S | **79.59%** | 82.67% | 82.83% | 76.41% | 76.45% | 76.45% |
| LambdaMART<sub>MAP</sub> | 78.36% | 82.19% | 82.29% | **77.67%** | **77.67%** | **77.67%** |
| ENR | 76.45% | 81.12% | 81.17% | 75.01% | 75.10% | 75.10% |

The best model for the CCLE data set (474 unique samples and 24 unique drugs) is ITNR with an NDCG@10 of 94.03% and an MRR@10 of 93.90%, with the ENR model close behind (NDCG@10 of 93.85% and MRR@10 of 93.64%). While DNN-S achieved the highest MRR among the baseline models, ITNR outperformed it by a relatively large margin. Moreover, although NDCG@5 of TNR-RankNet is higher than our model, it does not have comparable performance considering other evaluation metrics.

On the gCSI data set (357 unique samples and 16 unique drugs), ITNR demonstrates consistent performance improvement overall baseline models across all performance metrics and any of the chosen rank cutoffs. Notably, we observe a 2.6% performance improvement compared to the average of baseline models (i.e., 78.34%) in terms of NDCG@5. Moreover, ENR which performed really well on the CCLE data set demonstrated poor performance on the gCSI data set. TNR-RankNet and LambdaMART models achieved moderate performance and are the second-best methods. The models trained on gCSI did not generalize well. This was not surprising as gCSI had the smallest number of drugs and was thus prone to overfitting. Nevertheless, our method is able to maintain its high performance.

In general, the results for these two data sets indicate that ITNR significantly outperforms other competing methods. All models except ENR and LambdaMART are Deep Learning (DL) based models. Since DL models are complex and have many learnable parameters, they tend to overfit easier than traditional models. Due to this, DL algorithms perform better when trained on large amounts of data. To that end, we also conducted experiments on the GDSC data set (670 unique cell lines and 233 unique drugs) which is one of the largest public DRP data sets. Generally, DL models outperformed other traditional methods (i.e., ENR), since a model like ENR may not fully capture the structural information within drugs. Among transformer-based baselines, we observed that ITNR achieved the best performance, with 83.36% for NDCG@5 and 92.74% @25 of LambdaMART is higher than our model. However, this model is not even the second-best model considering other evaluation metrics. Among the competing models, TNR-RankNet outperformed other methods by a relatively large margin.

All in all, ITNR consistently outperforms all baseline methods across all metrics and data sets. In our experiment on three DRP data sets, TNR-RankNet and LambdaMART demonstrated reasonably good overall performance and they are the second-best methods. ITNR is not only able to push the most sensitive drugs to the top of the ranking list, but it can put them in the right order. Further, it can also generalize better and achieve superior performance on large DRP data sets like GDSC.

## 4. Discussion and Conclusion

Our study presented a novel transformer-based model, called Inversion Transformer-based Neural Ranking (ITNR), to predict cancer drug response. Our model used the well-known Transformer architecture to extract a better drugcell line representation. We also developed a novel loss function based on the concept of inversion and approximate permutation matrices. Our results suggest that the transformer network with multi-head attention is suitable for modeling the interactions of drug substructure and multiomics data. Extensive experimental results demonstrated that our model is more effective than the current state-of-the-art methods highlighting the predictive capability of our model and its potential translational value in personalized medicine. There are several directions for future work. In the current model, we use an approximated permutation matrix (i.e., NeuralSort) as the sorting operator. We can replace NeuralSort with another approximation of a sorting operator such as the Optimal Transport [7] and SoftSort [23]. Furthermore, the current model disregards the side effects and toxicity of drugs when predicting the best medication option. We can further optimize our recommendations by including the toxicity of drugs in our predictions.

## CRediT authorship contribution statement

**Shahabeddin Sotudian:** Conceptualization, Data curation, Formal analysis, Investigation, Methodology, Software, Validation, Visualization, Writing – Original Draft Preparation. **Ioannis Ch. Paschalidis:** Conceptualization, Formal analysis, Funding acquisition, Investigation, Methodology, Project administration, Resources, Supervision, Writing - review and editing.

## Appendix

### A. Hyper-parameter optimization

The list of hyper-parameters and their values for all ranking algorithms can be found in Table 2. In this table, *L* is the list length used for training (note that a list was either padded or sub-sampled to that length), *d_fc_* is the dimension of the linear projection, *N* is the number of encoder blocks, *d_h_* is the transformer hidden dimension, *H* is the number of attention heads, *D_r_* refers to the hidden dropout ratio, *B* is the batch size, *E* is the number of epochs, *ℓ*_2_ is the parameter of the *ℓ*_2_-norm penalty for the DNN-S model, *η* is the learning rate, *D_max_* is the maximum depth of a tree, *h_min_* is the minimum sum of the instance weight (Hessian) needed in a leaf, *N_T_* is the number of estimators, *α* is the regularization strength of the model, and *R*_*ℓ*_1__ controls the contribution of the *ℓ*_1_ and *ℓ*_2_ penalties in the ENR model. The details of hyper-parameter settings of all methods for the five folds can be found in Tables 3, 4, 5, and 6.

**Table 2.** The List of Hyper-parameters and Their Values.

| Algorithms | Parameters | Values |
| --- | --- | --- |
| All Transformer Models | $L$ | 120 |
| | $d_{fc}$ | 128, 256 |
| | $N$ | 2, 4 |
| | $d_h$ | 128, 512 |
| | $H$ | 2, 4 |
| DNN-S | $D_r$ | 0.1, 0.3, 0.5 |
| | $B$ | 25 |
| | $E$ | 50 |
| | $\ell_2$ | 0.01, 0.001, 0.0001 |
| LambdaMART_MAP | $\eta$ | 0.1, 0.01, 0.001 |
| | $D_{max}$ | 10, 100 |
| | $h_{min}$ | 1, 10, 1000 |
| | $N_T$ | 10, 100, 1000 |
| ENR | $\alpha$ | 0.01, 0.1, 1, 10, 100 |
| | $R_{\ell_1}$ | 0.2, 0.6, 0.8 |

**Table 3.** Best Parameters - Transformer Models

| Algorithms | Data Set | $L$ | $d_{fc}$ | $N$ | $d_h$ | $H$ |
| --- | --- | --- | --- | --- | --- | --- |
| Proposed Method | CCLE | 120 | 256 | 2 | 128 | 2 |
|  | GDSC | 120 | 256 | 2 | 128 | 2 |
|  | gCSI | 120 | 128 | 2 | 128 | 2 |
| TNR-NDCGLoss2++ | CCLE | 120 | 128 | 2 | 128 | 2 |
|  | GDSC | 120 | 256 | 2 | 128 | 2 |
|  | gCSI | 120 | 128 | 2 | 128 | 2 |
| TNR-ListMLE | CCLE | 120 | 256 | 2 | 128 | 2 |
|  | GDSC | 120 | 128 | 2 | 128 | 2 |
|  | gCSI | 120 | 128 | 2 | 128 | 2 |
| TNR-RankNet | CCLE | 120 | 128 | 2 | 128 | 2 |
|  | GDSC | 120 | 128 | 2 | 128 | 2 |
|  | gCSI | 120 | 128 | 2 | 128 | 2 |
| TNR-ApproxNDCG | CCLE | 120 | 256 | 2 | 128 | 2 |
|  | GDSC | 120 | 256 | 2 | 128 | 2 |
|  | gCSI | 120 | 128 | 2 | 128 | 2 |

**Table 4.** Best Parameters - DNN-S

| Data Sets | Folds | $B$ | $E$ | $D_r$ | $\ell_2$ |
| --- | --- | --- | --- | --- | --- |
| CCLE | Fold 1 | 25 | 50 | 0.3 | 0.0001 |
|  | Fold 2 | 25 | 50 | 0.5 | 0.01 |
|  | Fold 3 | 25 | 50 | 0.1 | 0.01 |
|  | Fold 4 | 25 | 50 | 0.3 | 0.001 |
|  | Fold 5 | 25 | 50 | 0.5 | 0.01 |
| GDSC | Fold 1 | 25 | 50 | 0.3 | 0.001 |
|  | Fold 2 | 25 | 50 | 0.5 | 0.01 |
|  | Fold 3 | 25 | 50 | 0.3 | 0.01 |
|  | Fold 4 | 25 | 50 | 0.3 | 0.001 |
|  | Fold 5 | 25 | 50 | 0.3 | 0.01 |
| gCSI | Fold 1 | 25 | 50 | 0.3 | 0.001 |
|  | Fold 2 | 25 | 50 | 0.3 | 0.001 |
|  | Fold 3 | 25 | 50 | 0.3 | 0.01 |
|  | Fold 4 | 25 | 50 | 0.3 | 0.01 |
|  | Fold 5 | 25 | 50 | 0.3 | 0.01 |

**Table 5.** Best Parameters - LambdaMART

| Data Sets | Folds | $\eta$ | $D_{max}$ | $N_T$ | $h_{min}$ |
| --- | --- | --- | --- | --- | --- |
| CCLE | Fold 1 | 0.1 | 10 | 100 | 1 |
|  | Fold 2 | 0.1 | 10 | 100 | 100 |
|  | Fold 3 | 0.1 | 100 | 100 | 1 |
|  | Fold 4 | 0.1 | 10 | 1000 | 10 |
|  | Fold 5 | 0.001 | 100 | 1000 | 1 |
| GDSC | Fold 1 | 0.1 | 10 | 1000 | 10 |
|  | Fold 2 | 0.001 | 10 | 100 | 1 |
|  | Fold 3 | 0.1 | 100 | 1000 | 10 |
|  | Fold 4 | 0.1 | 100 | 1000 | 1 |
|  | Fold 5 | 0.1 | 100 | 1000 | 1 |
| gCSI | Fold 1 | 0.01 | 10 | 1000 | 1 |
|  | Fold 2 | 0.1 | 10 | 1000 | 1 |
|  | Fold 3 | 0.001 | 10 | 1000 | 1 |
|  | Fold 4 | 0.1 | 10 | 1000 | 1 |
|  | Fold 5 | 0.1 | 10 | 1000 | 1 |

**Table 6.**
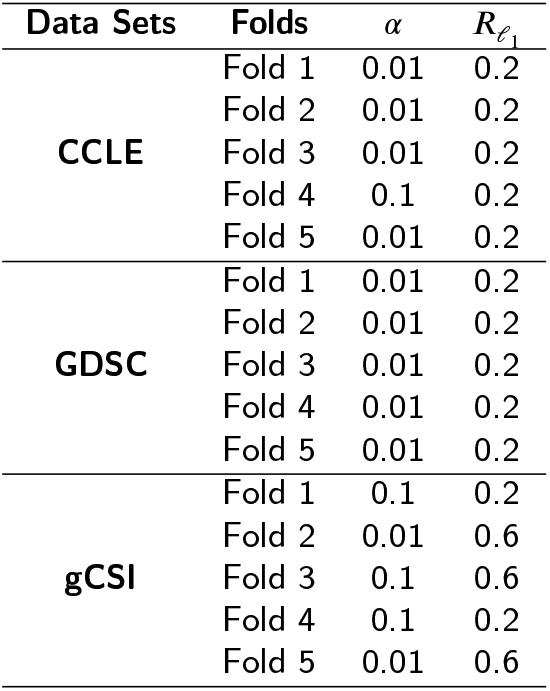
Best Parameters - ENR

